# Eco-friendly approach to indenodiazepinones from o-formylynones and o-phenylenediamines: Novel pharmacophoric constructs and their anti-bacterial activity

**DOI:** 10.64898/2026.01.01.697289

**Authors:** Bilal Ahmad Ganaie, Debapriya Mukherjee, Alagesan Balasubramani, Pallab Ghosh, Raju S Rajmani, Dipshikha Chakravortty, Goverdhan Mehta

**Author notes:** **Corresponding Authors:** Dipshikha Chakravortty - Department of Microbiology and Cell Biology, Indian Institute of Science, Bengaluru- 560012, India, Goverdhan Mehta - School of Chemistry, University of Hyderabad, Hyderabad 500 046, India; and. Equal contribution (Bilal Ahmad Ganaie, Debapriya Mukherjee).

## Abstract

We have conceptualised and executed an efficient, eco-friendly, one-flask, synthetic approach of general applicability to functionally enriched and potentially pharmacophoric tetracyclic indenodiazepinones through a base-mediated reaction cascade between a multifunctional *o*-formylynones and commercial *o*-phenylenediamines. The pertinence of this operationally simple, no-waste protocol, embodying many green and sustainable features, has been scoped with 26 variegated examples. This controlled one-pot assembly of novel tetracyclic indenodiazepinones with an unusual placement of an olefin through a reaction cascade involving Schiff base formation, intramolecular aza-Michael addition and a Mannich reaction between *o*-formylynones and *o*-phenylenediamines rapidly generates a diverse library poised for exploring their therapeutic potential and medicinal chemistry applications. The newly accessed indenodiazepinones exhibit promising antibacterial activities against many microorganisms and are non-cytotoxic even at higher concentrations. Interestingly, one of the compounds, **3v** did not develop resistance in *Salmonella* Typhimurium (STM) upon repeated exposure to sub-lethal doses up to the 160^th^ cycle, whereas resistance against clinically used antibacterial ciprofloxacin was attained at the 128^th^ cycle. These preliminary findings open avenues towards the development of novel antimicrobial agents with enhanced durability against drug resistance and these efforts are disclosed in this report.

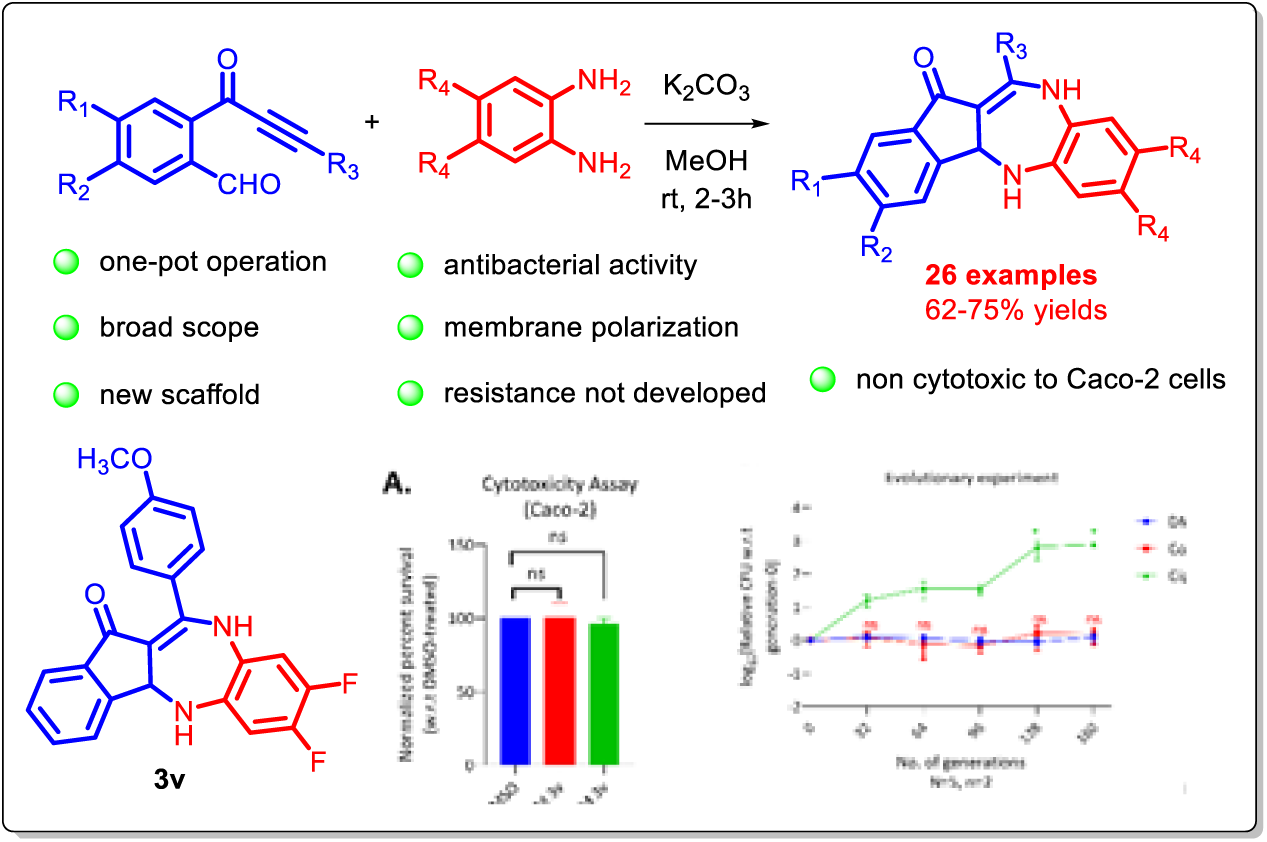

## INTRODUCTION

In the present times, with rising multiple global health concerns, the rapid advance of drug resistance has emerged as a major scientific challenge in the healthcare sector (1, 2) and WHO has highlighted “antimicrobial resistance (AMR) as a main threat to human beings.” (3) To thwart such forebodings and be future ready to overcome the lurking threat of AMR, a worldwide quest for new antimicrobial agents that are active against multidrug-resistant (MDR) microbes and function via alternative mode of action to stop, circumvent or delay its onset is an arena of utmost contemporary interest and poses a grave interdisciplinary and creativity challenge in drug discovery and clinical research and is garnering considerable traction. (4) While the need of the hour is an accelerated discovery and availability of new antibacterials, particularly those targeting MDR infections, the present reality in the drug discovery landscape is a slowdown with fewer FDA approvals in the past decade due to funding and profitability issues, despite its ascending importance and urgency in meeting the Sustainable Development Goals (SDGs). (5, 6) In this context, identification and access to new chemical space relevant to antimicrobial activity, as gleaned from clinically potent precedences, new target identification, and in silico projections are prerequisites.

The benzodiazepine moiety is a clinically well-established and successfully harnessed pharmacophoric scaffold in drug discovery programs across various therapeutic areas, including antimicrobial features. (7–10) A few representative bioactive benzodiazepine scaffold-based clinically used drugs, along with an example of an antimicrobial benzodiazepine derivative, are displayed in Figure 1. However, no benzodiazepine-based drug has been approved so far as an antimicrobial agent for clinical use. (11) Recognising this as a window of opportunity, we ventured to explore the potential of newly crafted, structurally modified and functionally enhanced benzodiazepines as antimicrobial agents. This mandated devising a *de novo*, versatile access to benzodiazepine frameworks, adhering to the key features of green and sustainable chemistry. (12) Due to their broad utility, numerous synthetic methods have been developed to access benzodiazepine frameworks. (13, 14) Literature reports indicate that the appropriate fusion of benzodiazepines with other pharmacophoric motifs often provides an entry to new chemical space with diverse and enhanced therapeutic profiles. Our ongoing interest (15–18) towards accessing complex indanone-fused carbo- and heterocycles from o-substituted ynones via cascade processes motivates us to devise a synthesis of indanone-fused benzodiazepine scaffolds from o-formylynones and o-phenylenediamines through a cascade process, involving Schiff base formation, intramolecular aza-Michael addition and Mannich reaction (Scheme 1), and assess their antimicrobial attributes against various pathogens.

**Figure 1.**
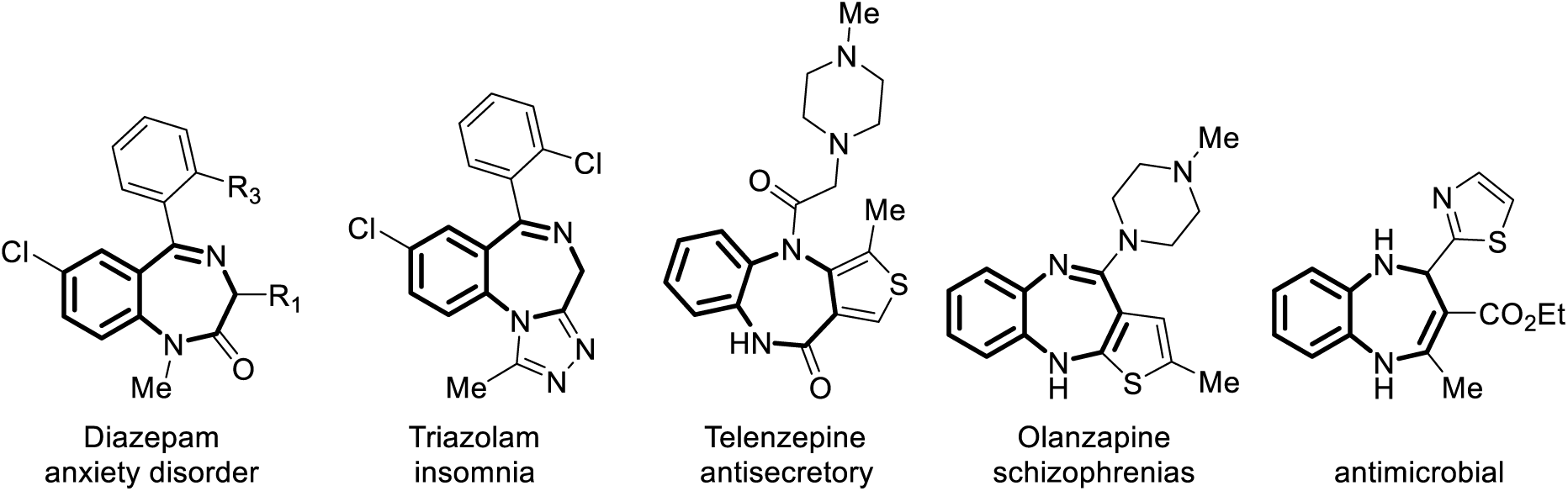
Representative bioactive benzodiazepine scaffolds

**Scheme 1.**
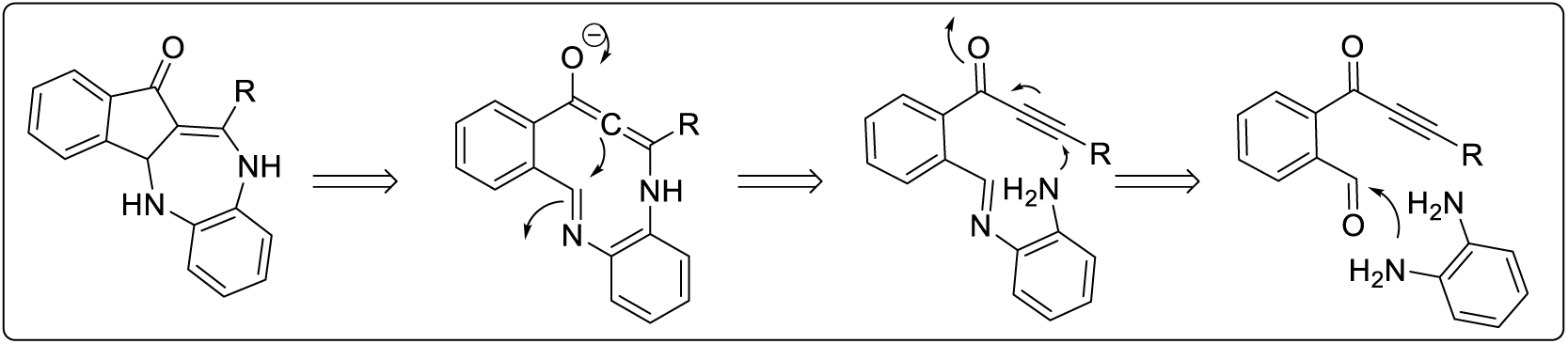
Conceptualisation of one-flak *de novo* access to functionally endowed benzodiazepines from o-formylynones and o-phenylenediamines.

## RESULTS AND DISCUSSION

As a pilot towards our exploratory foray to execute the conceptualisation of our benzodiazepine synthesis, o-formylynone **1a** was chosen for tandem engagement with o-phenylenediamine **2a** in the presence of a base. Pleasingly, the targeted product indanone fused benzodiazepinone **3a** was obtained and confirmed through its spectroscopic features (See SI). The reaction conditions were optimised by carrying out the reaction using K_2_CO_3_ as a base in MeOH solvent at ambient temperature to cleanly furnish the desired product **3a** (70%). For this type of exocyclically positioned double bond, containing indenodiazepinones, is a rarity. Having optimised conditions, it was imperative to scope this newly discovered indenodiazepinone synthesis methodology and demonstrate its generality employing diversely substituted *o*-formylynones and o-phenylenediamine. In this endeavour, a diversely substituted o-formyl ynones **1a**-**m**, including H, butyl, cyclopropyl, and electron-withdrawing and donating substituents on the pendant phenyl group of phenylacetylenes, were successfully utilized with o-phenylenediamine **2a** as a reacting partner to deliver corresponding indenodiazepinone **3a-l**. Furthermore, to enhance the binding with targets, substituted o-phenylenediamines **2b**-**2d** with –Me, Cl, and F were engaged with o-formylynones **1** to provide the corresponding indenodiazepinones **3m**-**3z** in good yields (72-82%), as detailed in Scheme 2. Further, the structures of indenodiazepinones **3** were secured based on their internally consistent spectral signatures and single-crystal X-ray diffraction analysis of representative examples, **3f**, **3j**, and **3n** (Scheme 2). As a scale-up experiment for the preparation of indenodiazepinone **3v** on a 10 mmol scale from **1f** and **2d** was uneventfully demonstrated (see Experimental Section).

**Scheme 2.**
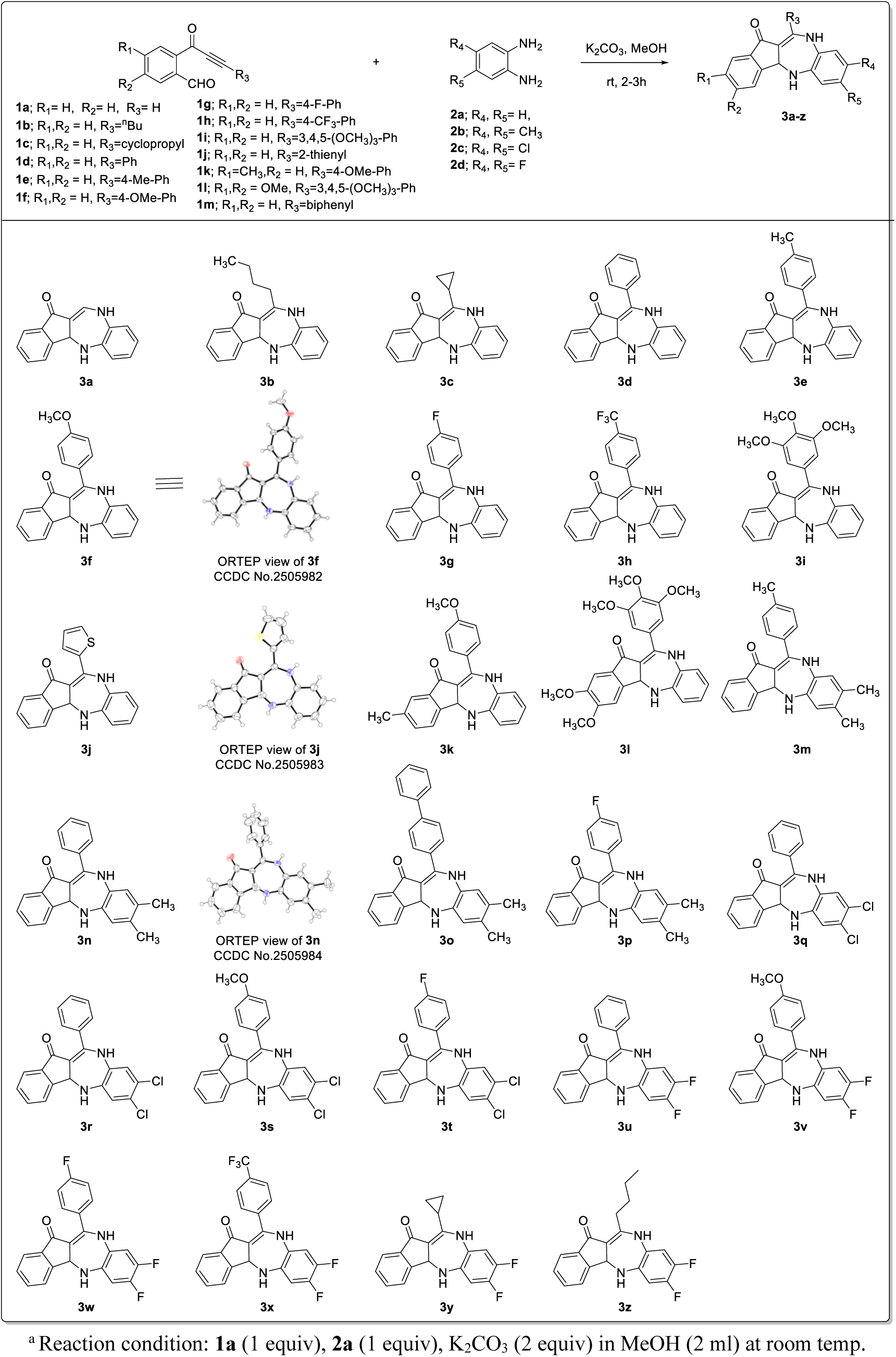
Scoping up of indenodiazepinones.

### Antimicrobial properties

This ready access to novel fused indenodiazepinone **3**, embodying pharmocophoric benzodiazepine and indanone moieties with multiple target binding potential, impelled us to undertake biological profiling of these new scaffolds. Initially, the characterization of the antimicrobial properties of compound **3** against the foodborne pathogen STM (19) was examined (See SI). A comparative analysis of antimicrobial efficacy demonstrated that while all derivatives exhibited antimicrobial activity, exposure to 30 µM of compounds **3v**, **3y**, and **3w** resulted in the lowest survival fraction compared to the solvent (DMSO) control. Ciprofloxacin, a commercially available antibiotic, served as a positive control, and at the same concentration, it exhibited greater bacterial killing than any of the tested compounds (Figure 2A) (20). Consequently, we concluded that our compounds had a bactericidal effect, though it was considerably weaker than ciprofloxacin at the same concentration. Compound **3v** was also found to be active against another notorious foodborne pathogen *Listeria* at a concentration of 7.3 µM (Figure S1A).

**Figure 2:**
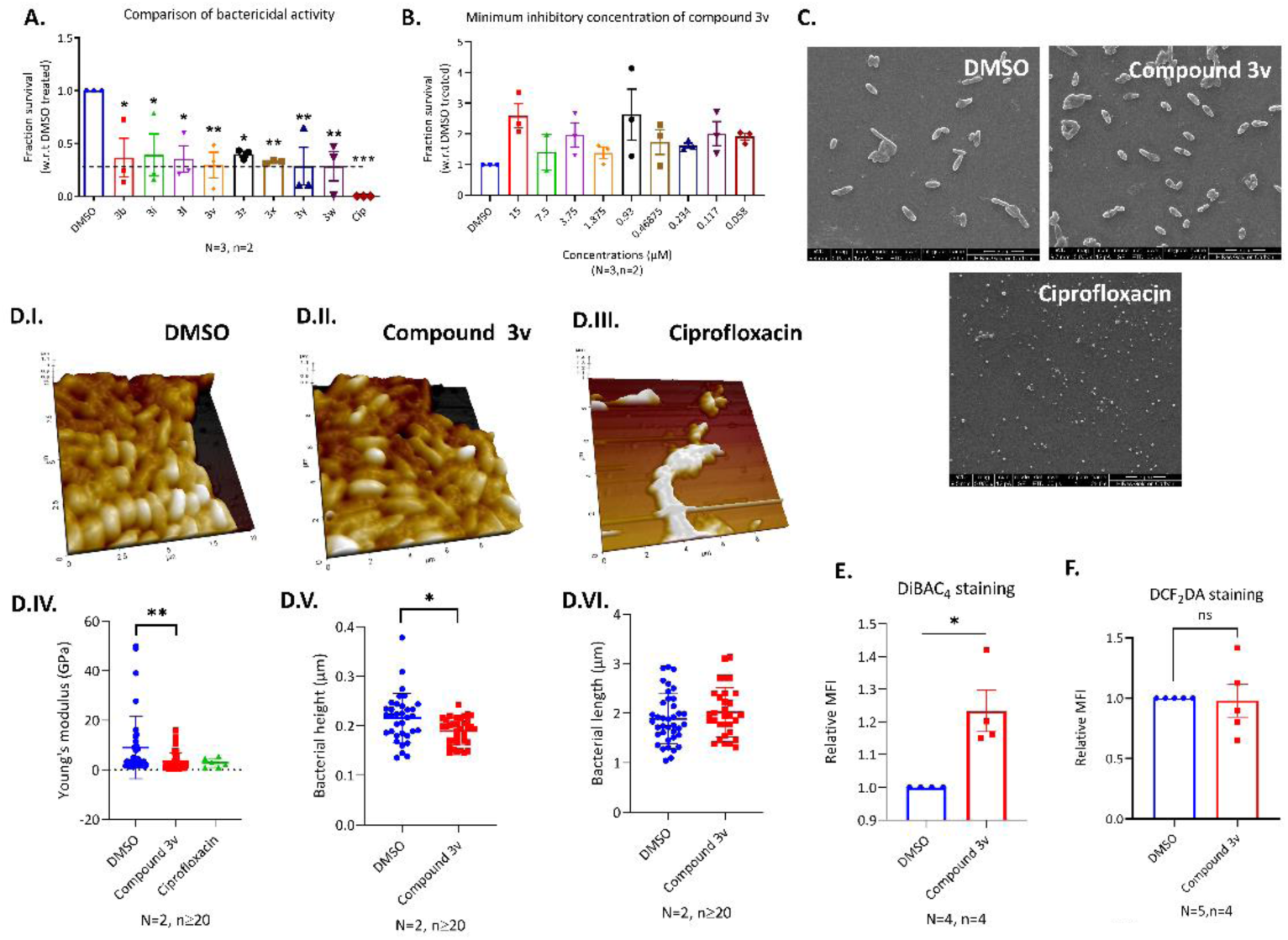
Derivatives of Compound 3 exhibit antimicrobial activity against STM, with membrane depolarization contributing to their mechanism of action. (A) Fractional survival of STM following exposure to 30 µM of Compound 3 derivatives and ciprofloxacin, compared to DMSO-treated controls. (B) Assessment of STM survival upon exposure to lower concentrations of 3v to determine its minimum inhibitory concentration. (C) Scanning Electron Microscopy and (D) Atomic Force Microscopy visualization of bacterial morphology after treatment with vehicle (DMSO), 30 µM Compound 3v, and ciprofloxacin, along with quantification of bacterial Young’s modulus, length, and height (D. IV–D. VI). (E) DiBAC_4_ and (F) DCF_2_DA staining of STM following exposure to Compound 3v. *p* < 0.05 (\**), p < 0.005 (**), p < 0.0005 (\*\*\**), *p* < 0.0001 (****), ns = non-significant. Statistical analyses were performed using an unpaired two-tailed Student’s *t*-test for comparisons between two groups and one-way ANOVA for comparisons involving more than two groups.

Further characterization focused on the antimicrobial activity of compound **3v**. Lower doses of compound **3v** did not exhibit a bactericidal effect on STM (Figure 2B), leading us to determine that 30 µM was the minimum concentration required for bactericidal activity. Thus, we used 30 µM for subsequent studies. Scanning electron microscopy (SEM) images revealed a reduction in bacterial size following treatment with compound **3v** (Figure 2C), which correlated with a shrunken and distorted surface morphology. Atomic Force Microscopy-assisted quantification of Young’s modulus further confirmed a reduction in bacterial stiffness upon exposure to compound **3v** (Figure 2D). Consistent with our SEM findings, we observed a significant decrease in bacterial height after treatment.

Given the observed changes in surface morphology, we performed DiBAC_4_ dye-assisted quantification of membrane polarization (Figure 2E) (21). An increase in relative MFI upon exposure to compound **3v** indicated membrane depolarization in STM, which may contribute to its antimicrobial activity (22). Unlike conventional antibiotics, treatment with compound **3v** did not lead to an increase in the relative MFI of DCF_2_DA, suggesting that its mechanism of bacterial killing was independent of oxidative stress induction (Figure 2F) (23).

### Compound 3v elicited antibacterial activities in *in-vivo* system and STM did not develop resistance to it till 160^th^ generation of exposure

To assess whether compound **3v** exhibited cytotoxic effects on eukaryotic cells, we conducted an MTT assay on the human colorectal carcinoma (Caco-2) cell line (Figure S1B). The results indicated that compound **3v** did not induce cytotoxicity in Caco-2 cells.

To evaluate the *in vivo* efficacy of compound **3v** in reducing bacterial load, we infected 6–8-week-old male Balb/c mice with 10^6^ CFU of STM and administered an intraperitoneal injection of 30 µM/gm-wt of compound **3v** or ciprofloxacin on days 2, 3, and 4 post-infection (Figure S1C). Upon euthanization at 5 dpi, no significant reduction in bacterial burden was observed in the blood, intestine, spleen, or liver of **3v**-treated mice compared to vehicle-treated controls. Similarly, ciprofloxacin at the same concentration was ineffective in reducing the bacterial load (Figure S1D).

To determine whether a higher dose of compound **3v** could impact bacterial burden, we tested 300 µM/gm-wt and 600 µM/gm-wt doses—10 and 20 times the initial concentration, respectively (Figure 3). An MTT assay on Caco-2 cells confirmed that these higher concentrations did not induce cytotoxicity (Figure 3A). Administration of 300 µM/gm-wt and 600 µM/gm-wt at the same dosing schedule resulted in a significant reduction in bacterial burden across the blood, intestine, spleen, and liver. The observed reduction was comparable to that achieved with the same doses of ciprofloxacin (Figure 3B, C. I–C. IV).

**Figure 3:**
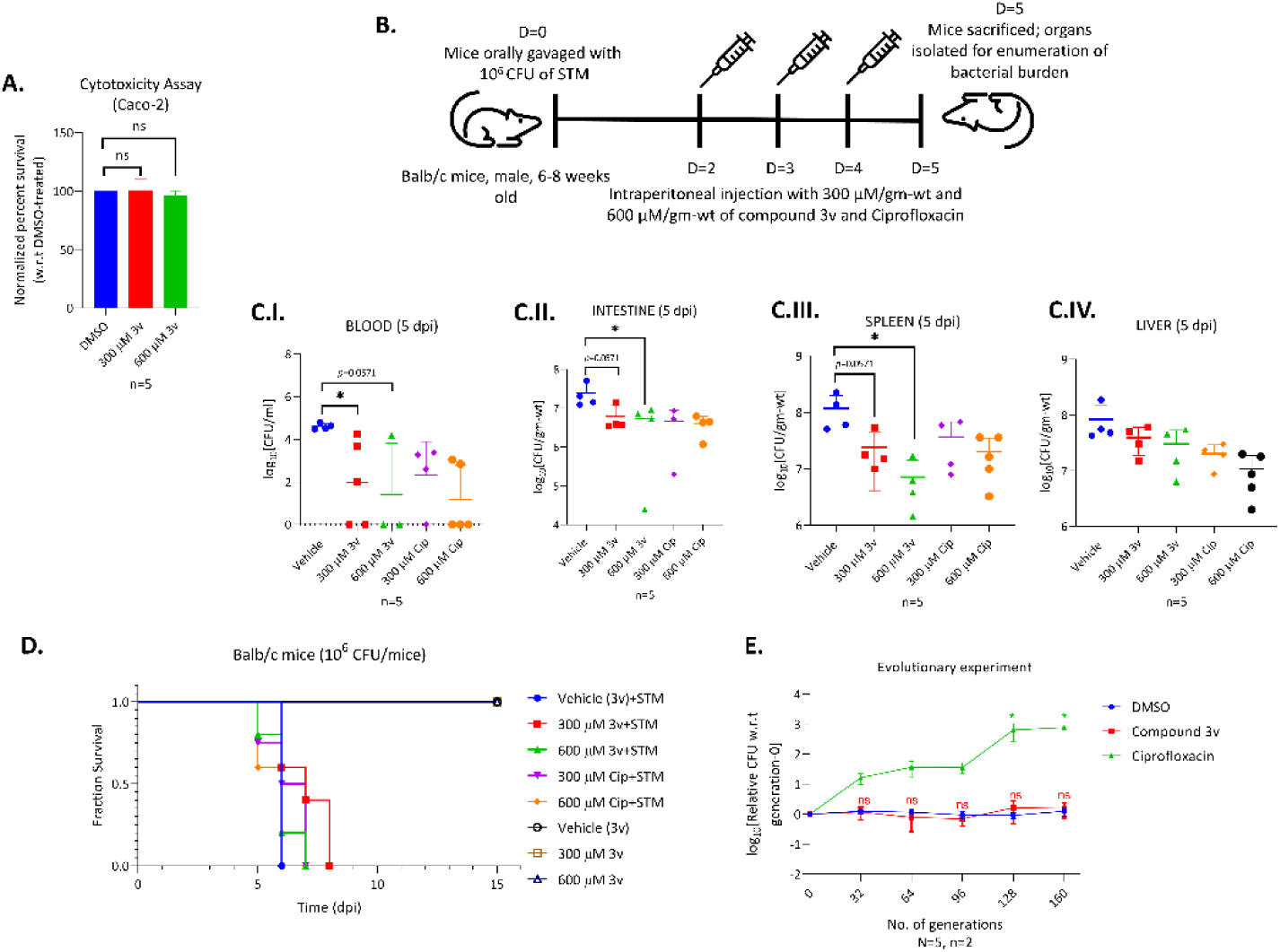
Compound 3v (300 and 600 µM/gm-wt) reduced bacterial load in STM-infected mice as effectively as ciprofloxacin, with no resistance development observed up to generation 160 *in-vitro*, unlike ciprofloxacin. (A) Cytotoxicity assay of Compound 3v on the human colorectal cell line Caco-2 at 300 µM and 600 µM. (B) Schematic representation of the infection and drug administration protocol in the Balb/c mouse infection model. (C) Bacterial burden in the blood (C.I), intestine (C.II), spleen (C.III), and liver (C.IV) five days post-infection. (D) Survival curve of Balb/c mice infected with 10⁶ CFU of STM and treated with 300 µM/gm-wt or 600 µM/gm-wt of Compound 3v and ciprofloxacin. (E) Line graphs showing log₁₀(relative CFU compared to generation 0) after exposure to lethal doses of Compound 3v and ciprofloxacin following repeated generations of sublethal dosing. *p* < 0.05 (\**), p < 0.005 (**), p < 0.0005 (\*\*\**), *p* < 0.0001 (****), ns = non-significant. Statistical analyses were conducted using one-way ANOVA for comparisons involving more than two groups and Mann-Whitney U-test for bacterial burden in animal experiments.

Hematoxylin and eosin (H&E) staining was performed on the fixed organ samples, and an expert pathologist conducted blinded pathological scoring (Figures S2 and S3). In the analysis of splenic histopathology, treatment with both 300 and 600 µM/gm-wt doses of the compounds significantly reduced the pathology scores compared to the group that received the vehicle only. These effects were comparable to those observed in the group treated with 600 µM/gm-wt ciprofloxacin (Figure S2). Similarly, in the liver tissue samples, administration of 600 µM/gm-wt of the compound was sufficient to lower the pathology score relative to the vehicle-treated group (Figure S3). Furthermore, treatment with both concentrations of compound **3v** successfully delayed mortality in STM-infected mice, similar to commercially available antibiotic ciprofloxacin (Figure 3D). Therefore, it can be concluded that compound **3v** remains active in the mouse model and is effective in reducing the Salmonella bacterial load, which is also reflected in the improvement of pathology scores.

Drug resistance remains a major challenge in drug discovery (24). To assess whether prolonged exposure to sublethal concentrations of compound **3v** could lead to resistance in STM, we exposed subsequent generations of the bacteria to sublethal doses of compound **3v** and ciprofloxacin. Bacterial samples from different generations were then exposed to a lethal concentration of **3v**. The CFU count remained unchanged compared to generation 0 up to the 160th generation for both compound **3v** and the vehicle control. However, ciprofloxacin-treated bacteria exhibited a drastic increase in relative CFU starting from generation 32, which became statistically significant at generations 128 and 160. This indicates that STM may develop resistance to ciprofloxacin after exposure to sublethal doses. However, exposure to sublethal doses of our compound **3v** did not lead to the emergence of resistance in STM. Furthermore, its effectiveness in treating STM infection in mice, comparable to conventional antibiotics, highlights its potential as a promising antimicrobial candidate.

## CONCLUSION

In summary, we have discovered a *de novo* synthesis of functionally enriched benzodiazepinones through a cascade reaction between o-formylynones and o-phenylenediamines through tandem Schiff base formation, intramolecular aza-Michael addition, Mannich reaction sequence. The new synthesis of indenodiazepinones has been scoped with 26 examples involving diversely substituted o-formylynones and o-phenylenediamines; all proceeding consistently in good to moderate yields with formation of three new σ-bonds (1C-C & 2C-N) and embellishment of functionalities in one-flask operation. These newly synthesised indenodiazepinones have been evaluated for their antibacterial activity against four microorganisms and among them *Salmonella* Typhimurium and *Listeria monocytogenes* were found to be more promising and were further explored. The compound **3v** was found to interfere with the membrane polarization of STM and that can be the plausible reason for the bacterial mortality. Importantly, STM did not attain resistance to compound **3v** after 160 repeated cycles. Compound **3v** was also found not to be toxic to normal cells, even at a higher dose of 300 mg/kg.

## Materials and Methods

### Bacterial strains

The *Salmonella enterica* subspecies *enterica* serovar Typhimurium strain 14028S, used in this study, was kindly provided by Professor Michael Hensel from the Max von Pettenkofer Institute for Hygiene and Medical Microbiology, Germany to DC Lab. *Listeria monocytogenes* ATCC 19112 was also utilized in this study. Bacterial strains were preserved as glycerol stocks at −80°C and revived on fresh Luria-Bertani (LB) agar plates (HiMedia) before the experiment commenced.

### Antimicrobial efficacy testing

To evaluate the antimicrobial activity of the compounds, a single colony from freshly revived stocks was inoculated into 5 mL of LB broth and incubated overnight at 37°C with shaking at 175 rpm. The overnight cultures were then subcultured at a 1:100 ratio and incubated under the same conditions for 6–8 hours to obtain cultures in the logarithmic phase. The OD₆₀₀ of the culture was adjusted to 0.3, after which the cultures were treated with 30 µM concentrations of the compounds and ciprofloxacin. Following 16 hours of incubation, the treated samples were plated on LB agar using the spread plating method. After an additional 16-hour incubation, the number of colonies was counted, and the fold decrease compared to the vehicle control was plotted to assess antimicrobial efficacy. A similar protocol was followed for bacterial cells treated with lower doses of compound 3v.

### Scanning Electron Microscopy

Logarithmic-phase STM cultures were incubated with 30 µM compound 3v and ciprofloxacin for 16 hours at 37°C with shaking at 175 rpm. After incubation, the cultures were drop-cast onto clean glass coverslips and air-dried. The samples were then fixed overnight at 4°C using 3.5% glutaraldehyde. The following day, they underwent a serial dehydration process with ethanol concentrations of 10%, 30%, 50%, 70%, 90%, and 100%. Subsequently, the samples were dried using a vacuum desiccator and subjected to gold sputtering prior to imaging. Image acquisition was performed using a Thermo Fischer XL-30 ESEM.

### Atomic Force Microscopy

The morphology and membrane elasticity of bacterial cells treated with the compound were analyzed using a modified protocol from Braga et al (25). The logarithmic phase culture of STM was treated with 30 µM of the compound 3v and incubated overnight at 37°C with shaking. Following incubation, 50 µL of the bacterial cultures were spotted onto coverslips and air-dried. The samples were subsequently fixed with 3.5% paraformaldehyde (PFA) for 10 minutes, washed with Milli-Q water, and dried using a vacuum desiccator. Finally, the bacterial cells were examined using an Atomic Force Microscope, and the Young’s Modulus was calculated with XEI software.

### DCF_2_DA staining

Logarithmic-phase STM cultures were treated with 30 µM of compound 3v and incubated overnight at 37°C under shaking conditions. Following incubation, 50 µL of each culture was transferred into wells of a 96-well plate, along with 50 µL of DCF_2_DA dye solution in PBS at a final concentration of 10 µM. The samples were then incubated at 37°C for 15 minutes. Fluorescence intensity was measured using a Tecan plate reader at an excitation/emission wavelength of 485/535 nm and normalized to bacterial cell absorbance at 600 nm (26).

### DiBAC_4_ staining

Logarithmic-phase STM was exposed to 30 µM of compound 3v and incubated overnight at 37°C under static conditions. Following incubation, 50 µL of the culture was transferred into each well of a 96-well plate, along with 50 µL of DiBAC_4_ dye solution in PBS at a final concentration of 1 µg/mL and incubated at 37°C for 15 minutes. Fluorescence intensity was then measured using a Tecan plate reader at an excitation/emission wavelength of 490/516 nm, with values normalized to the bacterial cell absorbance at 600 nm (26).

### Animal experiments

The animal experiments were approved by the Institutional Animal Ethics Clearance Committee (IAEC) at the Indian Institute of Science, Bangalore. All procedures strictly adhered to the guidelines established by the Committee for the Purpose of Control and Supervision of Experiments on Animals (CPCSEA). The ethical clearance number for this study is CAF/Ethics/973/2023.

To assess organ burden, 6–8-week-old male BALB/c mice were orally gavaged with 10⁶ CFU of STM on day 0. The compound, at concentrations of 30, 300, and 600 µM/g body weight, along with ciprofloxacin, was administered intraperitoneally on days 2, 3, and 4. On day 5 post-infection, the mice were euthanized, and samples of the liver, spleen, intestine, and blood were collected. The organs were homogenized, and bacterial load in each tissue was determined by spread plating on *Salmonella-Shigella* agar plates (27). For haematoxylin and eosin staining, liver and spleen tissues were fixed and preserved in 3.5% paraformaldehyde. The staining procedure was outsourced, and histopathological scoring was performed in a blinded manner by a qualified pathologist.

A similar infection and drug administration protocol was followed for the survival curve analysis, with mice being monitored until day 15 to assess mortality.

### Resistance Development

The protocol for the adaptive laboratory evolution of (STM) exposed to a sub-lethal concentration of the compound and the antibiotic was adapted from Ghoshal *et al* (28). Logarithmic phase of STM was treated with 15 µM of the compound (sub-MIC dose) and 0.018 µM ciprofloxacin (sub-MIC dose) and incubated overnight at 37°C under static conditions. Serial passaging of the treated bacterial cells was performed in fresh media while maintaining the specified treatment concentrations for a total of 160 generations. Glycerol stocks were prepared at generations 0 (before treatment initiation), 32, 64, 128, and 160.

To assess resistance development, 20 µL of bacterial cultures from each generation was inoculated from frozen glycerol stocks into 5 mL of fresh LB broth and incubated overnight at 37°C with shaking. These cultures were then subcultured in fresh media and incubated at 37°C for 5 hours under shaking conditions to reach the logarithmic phase. Bacterial cells were subsequently treated with 30 µM (MIC dose) of the compound and 0.037 µM ciprofloxacin (MIC dose) and incubated overnight at 37°C under static conditions. The CFU/mL of STM at each generation, following treatment with 30 µM of the compound and 0.037 µM ciprofloxacin, was determined by bacterial spread plating. A log₁₀(Relative CFU w.r.t generation-0) was plotted to analyse resistance development.

### Statistical Analysis

The number of biological and technical replicates for each experiment is specified in the figure panels. Statistical analyses were conducted using an unpaired two-tailed Student’s *t*-test or one-way ANOVA, as indicated in the figure legends. A *p*-value of less than 0.05 was considered statistically significant. Results are presented as mean ± SEM. All the graphs and statistical tests were generated using GraphPad Prism 8.4.3 (686).

## ASSOCIATED CONTENT

## Supporting Information

The Supporting Information is available free of charge on the ACS Publications website. Crystallographic data and copies of NMR spectra for compound **3a-z** (PDF)

## Accession Codes

CCDC 2505982-2505984 contains the supplementary crystallographic data for this paper. These data can be obtained free of charge via www.ccdc.cam.ac.uk/data_request/cif, or by emailing, or by contacting The Cambridge Crystallographic Data Centre, 12 Union Road, Cambridge CB2 1EZ, UK; fax: +44 1223 336033.

## AUTHOR INFORMATION

## Notes

The authors declare no competing financial interest.

## Acknowledgments

B.A.G acknowledges SERB for the award of the National Postdoctoral Fellowship (NPDF). G.M. acknowledges Dr. Reddy’s Laboratories for the award of Dr. Kallam Anji Reddy Chair Professorship. DM acknowledges the IISc fellowship provided by MHRD, Government of India. PG appreciates the DBT-JRF fellowship. DC expresses gratitude for the ASTRA Chair Professorship grant from IISc and the TATA Innovation Fellowship grant. This research was financially supported by the DAE SRC fellowship (DAE00195) and the DBT-IISc partnership umbrella program for advanced research in biological sciences and bioengineering. The authors also acknowledge technical support from ICMR (Centre for Advanced Study in Molecular Medicine), DST (FIST), and UGC (Special Assistance). Additionally, they extend their thanks to IISc and DST-SERB for financial assistance through the research grant EMR/2017/001016.

## Supplementary figures and figure legends

**Figure S1:**
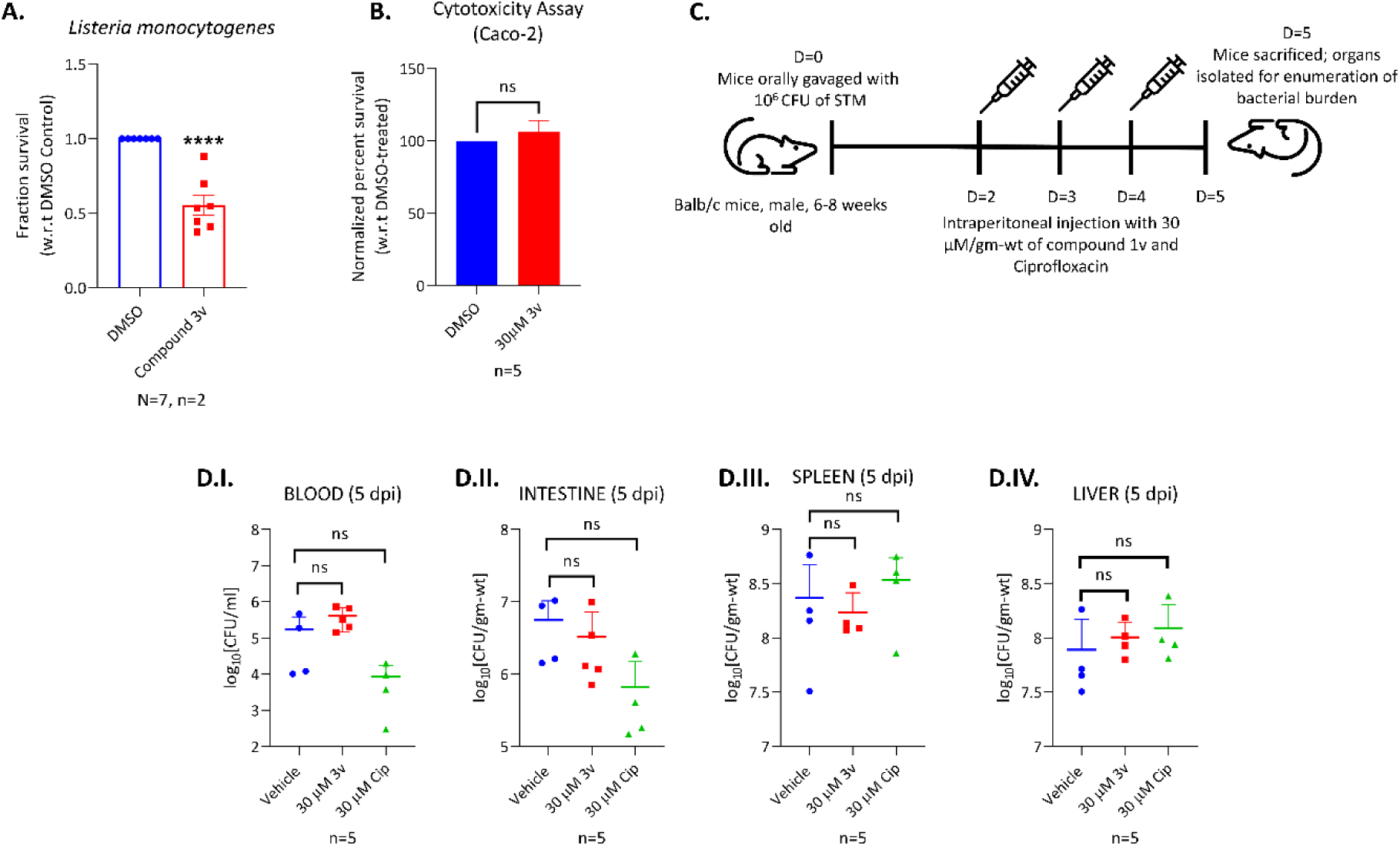
Compound 3v at 30 µM/gm-wt was ineffective in reducing bacterial load in STM-infected mice. (A) Bar graph showing the fraction survival of *Listeria monocytogenes* after exposure to 7.3 µM of Compound 3v. (B) Cytotoxicity assay of Compound 3v on the human colorectal cell line Caco-2 at 30 µM. (C) Schematic illustration of the infection and drug administration protocol in the Balb/c mouse infection model. (D) Bacterial burden in the blood (D. I), intestine (D. II), spleen (D.III), and liver (D.IV) five days post-infection. *p* < 0.05 (\**), p < 0.005 (**), p < 0.0005 (\*\*\**), *p* < 0.0001 (****), ns = non-significant. Statistical analyses were performed using one-way ANOVA for comparisons involving more than two groups and the Mann-Whitney U-test for bacterial burden in animal experiments.

**Figure S2:**
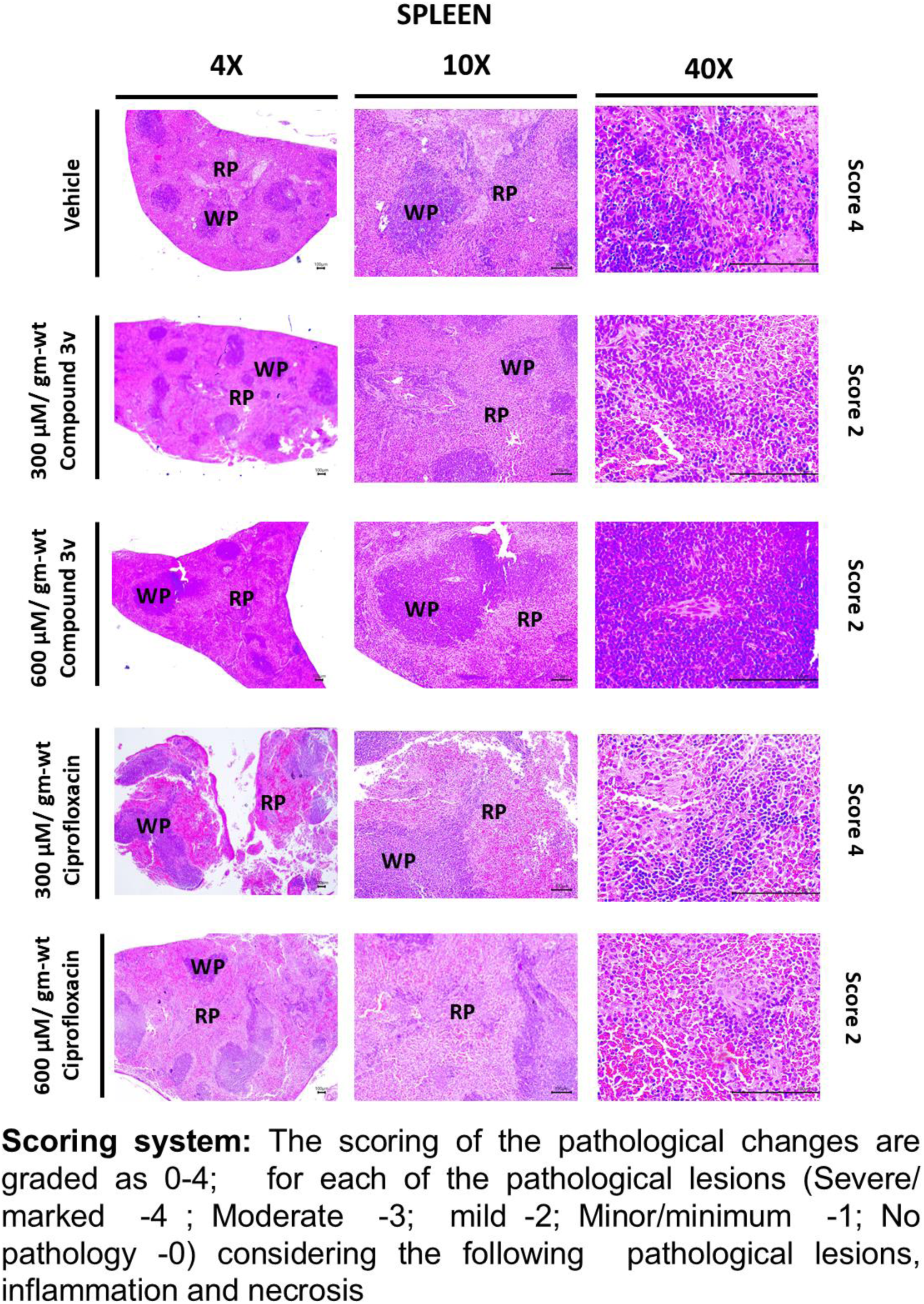
Administration of the compound 3v at both 300 and 600 µM/gm-wt doses led to a significant decrease in pathology scores when compared to the vehicle-treated group. Representative haematoxylin-eosin-stained spleen section images illustrating the effects of compound 3v and ciprofloxacin treatments at 300 and 600 µM/gm-wt in *Salmonella*-infected mice on day 5 post-infection, captured at 4X, 10X, and 40X magnifications.

**Figure S3:**
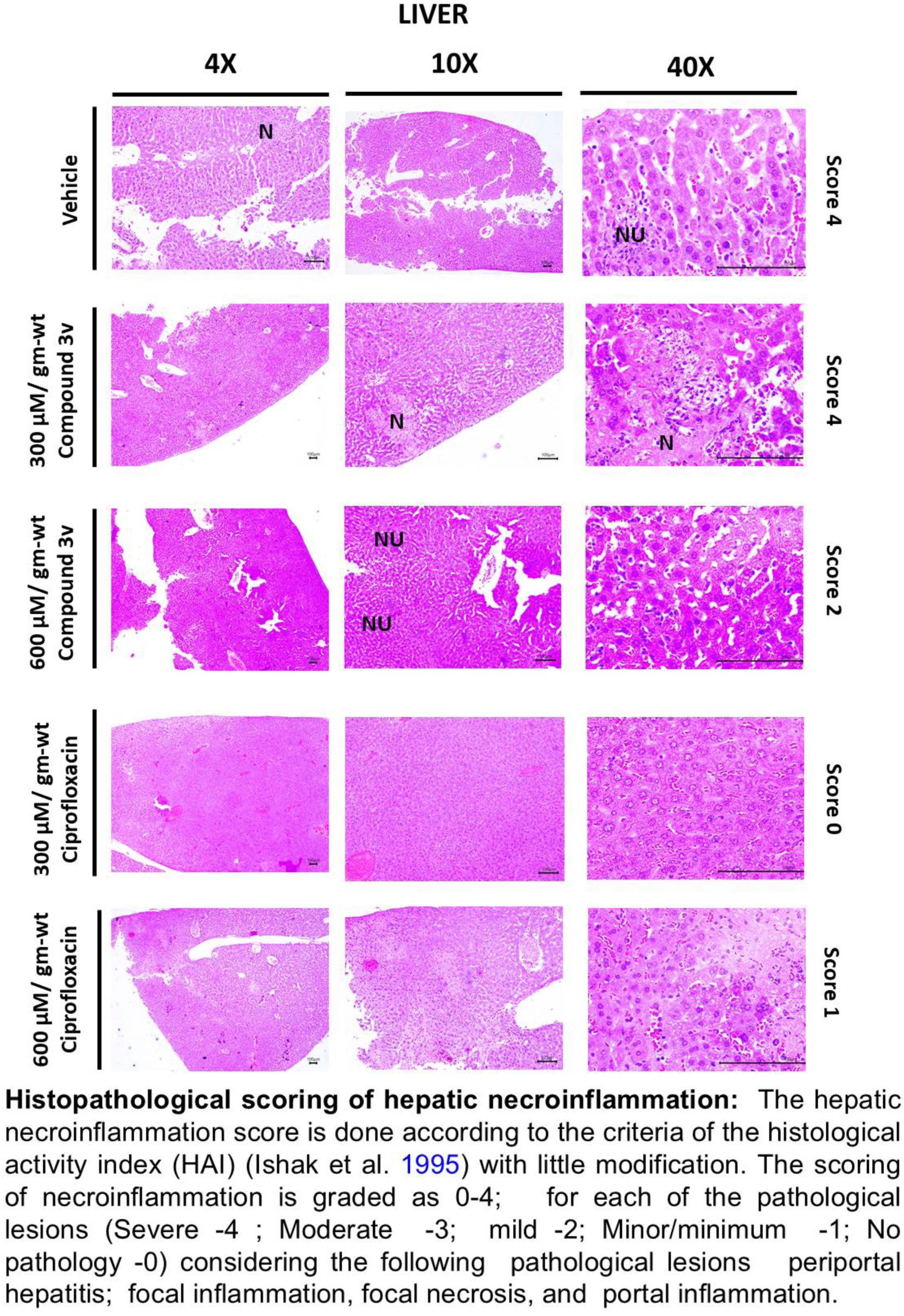
Treatment with compound 3v at a dose of 600 µM/gm-wt resulted in a reduction in liver pathology scores compared to the vehicle-treated group. Representative haematoxylin-eosin-stained spleen section images depict the effects of compound 3v and ciprofloxacin administered at 300 and 600 µM/gm-wt in *Salmonella*-infected mice on day 5 post-infection, captured at 4X, 10X, and 40X magnifications. N= Necrosis, NU= Neutrophil accumulation.

## References

1. Kurt Yilmaz N, Schiffer CA. Introduction: Drug Resistance. Chem Rev. 2021;121(6):3235–7.

2. Saha M, Sarkar A. Review on Multiple Facets of Drug Resistance: A Rising Challenge in the 21st Century. J Xenobiot. 2021;11(4):197–214.

3. WHO. https://www.who.int/news-room/fact-sheets/detail/antimicrobial-resistance.

4. Annunziato G. Strategies to Overcome Antimicrobial Resistance (AMR) Making Use of Non-Essential Target Inhibitors: A Review. Int J Mol Sci. 2019;20(23).

5. Qiao, W.; Wang, L.; Luo, Y.; Yang, T. Synthetic approaches and therapeutic applications of FDA-approved antibacterial agents: A comprehensive review from 2003 to 2023. Eur. J. Med. Chem. 2025, 285, 117267–117291

6. https://sdgs.un.org/goals

7. Tolu-Bolaji OO, Sojinu SO, Okedere AP, Ajani OO. A review on the chemistry and pharmacological properties of benzodiazepine motifs in drug design. Arab Journal of Basic and Applied Sciences. 2022;29(1):287–306.

8. Edinoff AN, Nix CA, Odisho AS, Babin CP, Derouen AG, Lutfallah SC, et al. Novel Designer Benzodiazepines: Comprehensive Review of Evolving Clinical and Adverse Effects. Neurol Int. 2022;14(3):648–63.

9. Manchester KR, Lomas EC, Waters L, Dempsey FC, Maskell PD. The emergence of new psychoactive substance (NPS) benzodiazepines: A review. Drug Test Anal. 2018;10(1):37–53.

10. Nielsen S. Benzodiazepines. Non-medical and illicit use of psychoactive drugs: Springer; 2015. p. 141–59.

11. https://www.ncbi.nlm.nih.gov/books/NBK470159

12. https://www.unep.org/topics/chemicals-and-pollution-action/circularity-sectors/green-and-sustainable-chemistry

13. Farhid, H.; Khodkari, V.; Nazeri, M. T.; Javanbakht, S.; Shaabani, A. Multicomponent reactions as a potent tool for the synthesis of benzodiazepines. Org. Biomol. Chem. 2021, 19 (15), 3318–3358

14. Arora, N.; Dhiman, P.; Kumar, S.; Singh, G.; Monga, V. Recent Advances in Synthesis and Medicinal Chemistry of Benzodiazepines. Bioorg. Chem. 2020, 97, 103668.

15. Gunnam A, Balasubramani A, Mehta G. Recursive Anion-Triggered Tandem Reactions of ortho-Bis-ynones: Tunable Synthesis of 1-Indenones and Cyclopenta[a]inden-8(2H)-ones. J Org Chem. 2022;87(6):4376–84.

16. Balasubramani, A.; Gunnam, A.; Mehta, G. In situ Generated Ammonia Mediates Deep Restructuring of o-Bis-ynones Through a Cascade Process: One-pot synthesis of 2-Azafluorenones. J. Org. Chem. 2022, 87, 10138–10145

17. Balasubramani A, Mehta G. One-pot synthesis of functionally enriched benzo [b] fluorenones: An eco-friendly embedment of diverse 1-Indanones into o-Bis-Ynones. The Journal of Organic Chemistry. 2023;88(2):933–43.

18. Ganaie BA, Balasubramani A, Bhat BA, Mehta G. Indeno-Annulation of o-Formyl-Ynones, o-Bis-Ynones, and p-Bis-o-Formyl-Ynones with Dimethyl Acetone-1,3-Dicarboxylate: One Flask Cascade Synthesis of Functionally Endowed 9-Fluorenols and Indeno[1,2-b]fluorenols. J Org Chem. 2023;88(16):11637–49.

19. Liu X, Liu Y, Zhao X, Li X, Yao T, Liu R, et al. Salmonella enterica serovar Typhimurium remodels mitochondrial dynamics of macrophages via the T3SS effector SipA to promote intracellular proliferation. Gut Microbes. 2024;16(1):2316932.

20. Shariati A, Arshadi M, Khosrojerdi MA, Abedinzadeh M, Ganjalishahi M, Maleki A, et al. The resistance mechanisms of bacteria against ciprofloxacin and new approaches for enhancing the efficacy of this antibiotic. Front Public Health. 2022;10:1025633.

21. Mitchell SJ, Pardo-Pastor C, Zangle TA, Rosenblatt J. Voltage-dependent volume regulation controls epithelial cell extrusion and morphology. bioRxiv. 2023.

22. Molina-Hernandez JB, Aceto A, Bucciarelli T, Paludi D, Valbonetti L, Zilli K, et al. The membrane depolarization and increase intracellular calcium level produced by silver nanoclusters are responsible for bacterial death. Sci Rep. 2021;11(1):21557.

23. Dwyer DJ, Belenky PA, Yang JH, MacDonald IC, Martell JD, Takahashi N, et al. Antibiotics induce redox-related physiological alterations as part of their lethality. Proc Natl Acad Sci U S A. 2014;111(20):E2100–9.

24. Metlay JP, Powers JH, Dudley MN, Christiansen K, Finch RG. Antimicrobial drug resistance, regulation, and research. Emerg Infect Dis. 2006;12(2):183–90.

25. Braga PC, Ricci D. Atomic force microscopy: application to investigation of Escherichia coli morphology before and after exposure to cefodizime. Antimicrob Agents Chemother. 1998;42(1):18–22.

26. Chowdhury AR, Mukherjee D, Singh AK, Chakravortty D. Loss of outer membrane protein A (OmpA) impairs the survival of Salmonella Typhimurium by inducing membrane damage in the presence of ceftazidime and meropenem. J Antimicrob Chemother. 2022;77(12):3376–89.

27. Nair AV, Singh A, Rajmani RS, Chakravortty D. Salmonella Typhimurium employs spermidine to exert protection against ROS-mediated cytotoxicity and rewires host polyamine metabolism to ameliorate its survival in macrophages. Redox Biol. 2024;72:103151.

28. Ghoshal M, Bechtel TD, Gibbons JG, McLandsborough L. Adaptive laboratory evolution of Salmonella enterica in acid stress. Front Microbiol. 2023;14:1285421.

